# Fitness, Physical Activity, and Cardiovascular Disease: Longitudinal and Genetic Analyses in the UK Biobank Study

**DOI:** 10.1101/136192

**Authors:** Emmi Tikkanen, Stefan Gustafsson, Erik Ingelsson

## Abstract

**Background:** Exercise is inversely related with cardiovascular disease (CVD), but large-scale studies of incident CVD events are lacking. Moreover, little is known about genetic determinants of fitness and physical activity, and modifiable effects of exercise in individuals with elevated genetic risk of CVD. Finally, causal analyses of exercise traits are limited.

**Methods:** We estimated associations of grip strength, physical activity, and cardiorespiratory fitness with CVD and all-cause death in up to 502,635 individuals from the UK Biobank. We also examined these associations in individuals with different genetic burden on coronary heart disease (CHD) and atrial fibrillation (AF). Finally, we performed genome-wide association study (GWAS) of grip strength and physical activity, as well as Mendelian randomization analysis to assess the causal role of grip strength in CHD.

**Findings:** Grip strength, physical activity, and cardiorespiratory fitness showed strong inverse associations with incident cardiovascular events and all-cause death (for composite CVD; HR, 0.78, 95% CI, 0.77-0.80; HR, 0.94, 95% CI, 0.93-0.95, and HR, 0.67, 95% CI, 0.63-0.71, per SD change, respectively). We observed stronger associations of grip strength with CHD and AF for individuals in the lowest tertile of genetic risk (P_interaction_ = 0.006, P_interaction_ = 0.03, respectively), but the inverse associations were present in each category of genetic risk. We report 27 novel genetic loci associated with grip strength and 2 loci with physical activity, with the strongest associations in *FTO* (rs56094641, P=3.8×10^-24^) and *SMIM2* (rs9316077, P=1.4×10^-8^), respectively. By use of Mendelian randomization, we provide evidence that grip strength is causally related to CHD.

**Interpretation:** Maintaining physical strength is likely to prevent future cardiovascular events, also in individuals with elevated genetic risk for CVD.

**Funding:** National Institutes of Health (1 R01 HL135313-01), Knut and Alice Wallenberg Foundation (2013.0126), and the Finnish Cultural Foundation.

## Introduction

Cardiovascular disease (CVD) is a major public health issue and societal burden worldwide. Exercise has been highlighted as a cost-effective prevention strategy for CVD; it improves cardiorespiratory fitness (CRF) and muscular strength, which both have shown to be inversely associated with future CVD events in population-based studies^1,2^. However, fitness and physical activity are hard to measure accurately and consistently on a large scale; and thus, observational analyses prospectively relating fitness and physical activity with new-onset CVD among healthy individuals have typically been limited to smaller study samples. Moreover, so far, little progress has been made in disentangling the genetic determinants of fitness and physical activity. Such knowledge can provide important insights to physiological mechanisms related to exercise, allow for studies of gene-environment interactions, and of causality utilizing Mendelian randomization (MR) methodology.

MR is a useful method for evaluating causality of associations between predisposing risk factors and disease. Utilizing genetic data, MR analysis is by design unaffected by reverse causation, which is a common problem in observational epidemiological studies. Further, under certain assumptions, MR methods also avoid confounding, another major problem in observational studies. MR has been increasingly utilized in the past few years after genome-wide association studies (GWAS) have uncovered genetic variants robustly associated with various phenotypes. For example, MR studies have given the support for causal role of LDL cholesterol, triglycerides, and obesity, but not of HDL cholesterol, in CVD^3-5^.

In this paper, we analyzed objective and subjective measures of fitness and physical activity together information of CVD risk factors and genomics in relation to prospective CVD disease events and all-cause death in up to 502,635 individuals from the UK Biobank. We had three aims: 1) to study the associations of fitness and physical activity in relation to new-onset CVD and all-cause death in a large observational study and to assess whether these associations were modified by genetic risk; 2) to explore genetic determinants of measures of fitness and physical activity; and 3) to assess the causal role of fitness and physical activity in CVD.

## Material and Methods

UK Biobank is a major resource aiming to improve human health, and the prevention, diagnosis, and treatment of chronic diseases^6^. In this study, the four exposures of interest were grip strength, objective and subjective physical activity, and CRF **(Figure S1).** Grip strength was measured using a Jamar J00105 hydraulic hand dynamometer. We calculated relative grip strength as an average of measurements of right and left hand divided by weight (expressed as kg)^7,8^. Physical activity was assessed with a short form IPAQ questionnaire (“IPAQ-PA”)^9^ and with Axivity AX3 wrist-worn triaxial accelerometer^10^. CRF was assessed with net oxygen consumption (VO_2_), calculated from individuals’ body weight and maximum workload during the cycle ergometry on a stationary bike (eBike, Firmware v1.7) using the equation VO2=7+10.8(workload)/weight^11^. The CVD outcomes (coronary heart disease [CHD], ischemic stroke, hemorrhagic stroke, heart failure, atrial fibrillation [AF], and combined CVD) were defined using in-patient hospital and death registry data that have been linked to the UK Biobank (ICD codes in the **Supplementary Appendix).** All-cause death was defined from the death registry.

We estimated associations of grip strength, IPAQ-PA, and CRF on CVD events in up to 484,918 individuals using Cox proportional hazards models free from CVD at baseline. In secondary analyses, we also analyzed associations with all-cause death in up to 502,635 individuals. Accelerometer data was used for all-cause death analysis only, due to too short follow-up with hospital registry data. For each endpoint, we ran three sets of multivariable-adjusted models: a) adjusting for age, gender, and region of the UK Biobank assessment center; b) additional adjustment for possible confounders (ethnicity, BMI, smoking, lipid medication, systolic blood pressure, diabetes, height, and Townsend index); and c) adjusting for IPAQ-PA and/or grip strength in addition to those in b). We also examined the effects stratified by tertiles of a weighted genetic risk scores (GRS) for CHD and AF, and tested for interactions between the measures of fitness and physical activity, and the GRS. The genetic markers and the weights were selected from the largest published GWAS of CHD and AF (CARDIoGRAMplusC4D^12^ and AFgen^13^). Models including genetic data were further adjusted for genotype array and principal components.

We then performed GWAS for grip strength and IPAQ-PA to explore genetic determinants of fitness and physical activity. The discovery analyses were conducted in 80,000 randomly sampled individuals of European descent, and 40,285 were used for replication. The models were adjusted for age, gender, genotype array, and 15 principal components. Analyses were conducted with PLINK^14^ (version 1.9) by use of linear regression assuming an additive model for association between phenotypes and genotype dosages. We also conducted a pooled analysis for the whole sample to reveal additional loci associated with fitness and physical activity. For functional annotation, we used the GTEx portal^15^ and DEPICT^16^.

Finally, we performed two-sample MR^17, 18^ by using the genetic variants from the GWAS of grip strength as an instrumental variable (IV) and publicly available GWAS data for CHD (from CARDIoGRAMplusC4D^12^) as outcomes. Analyses were conducted with the R package *TwoSampleMR*^19^. Power for MR analyses was estimated with an online tool created by Burgess^20^.

A detailed description of material and methods can be found in the **Supplementary Appendix.**

## Role of the funding source

The sponsors had no role in the design of the study; collection, analysis, and interpretation of the data; or writing the report. The corresponding author had full access to the data and responsibility for the decision to submit for publication.

## Results

The study characteristics are shown in Table 1. Mean age at baseline was 56.5 years (SD, 8.1 years) and 54% of subjects were females. During follow-up (median 6.1 years; interquartile range, 5.4-6.7 years; 2,913,057 person-years at risk), 15,647 incident CVD cases occurred in participants free from the disease at baseline (8,176 CHD, 2,113 ischemic stroke, 1,012 hemorrhagic stroke, 928 heart failure, and 4,449 AF events).

**Table 1.** Baseline characteristics of the UK Biobank (N=502,635)

| **Table 1. Baseline characteristics of the UK Biobank (N=502,635)** |  |  |
| --- | --- | --- |
| Gender |  |  |
|  | Females | 273,465 (54%) |
|  | Males | 229,173 (46%) |
| Baseline age, years |  | 56.5 (8.1) |
| Ethnicity* |  |  |
|  | White | 475,378 (94.6%) |
|  | Black | 8,152 (1.6%) |
|  | Asian | 11,534 (2.3%) |
|  | Mixed | 7,574 (1.5%) |
| Smoking status* |  |  |
|  | Never | 275,221 (54.8%) |
|  | Previous | 174,129 (34.6%) |
|  | Current | 53,288 (10.6%) |
| Body mass index*, kg/m<sup>2</sup> |  | 27.4 (4.8) |
| Blood pressure*, mmHg |  |  |
|  | Systolic | 139.8 (19.7) |
|  | Diastolic | 82.3 (10.7) |
| Diabetes* |  | 26,587 (5.3%) |
| Lipid medication |  | 82,369 (16.4%) |
| Cardiovascular disease at baseline |  | 17,017 (3.4%) |
| Grip strength*, kg |  | 0.40 (0.13) |
| Physical activity*, MET-hours / week |  | 43.8 (43.7) |
| Cardiorespiratory fitness, 7+10.8(watts)/kg |  | 18.5 (3.3) |
| Data are mean (SD) or N (%). |  |  |
| * Missing values of the variable were imputed with predictive mean matching <sup>39</sup>. |  |  |

### Observational analyses

The results from observational analyses are shown in Table 2. We found strong inverse associations between grip strength and all outcomes (hazard ratios [HR] between 0.48 for heart failure and 0.91 for hemorrhagic stroke) in our age, gender, and region-adjusted models (models a). Effects were slightly attenuated when adjusting for confounding factors, but for all outcomes except AF, still highly significant. Grip strength was associated with these endpoints also after adjusting for IPAQ-PA.

**Table 2.** Incidence and hazard ratios for CVD endpoints by fitness and physical activity.

|  |  |  |  |  |  |  |  |  |  |  |
| --- | --- | --- | --- | --- | --- | --- | --- | --- | --- | --- |
|  |  | CVD |  |  | CHD |  |  | Ischemic stroke |  |  |
|  | Model | N events | HR (95 % CI) | P-value | N events | HR (95 % CI) | P-value | N events | HR (95 % CI) | P-value |
| Grip strength | 1 | 15,647 | 0.78 ( 0.77 , 0.80 ) | <0.0001 | 8,176 | 0.79 ( 0.77 , 0.81 ) | <0.0001 | 2,113 | 0.71 ( 0.67 , 0.75 ) | <0.0001 |
|  | 2 | 15,647 | 0.88 ( 0.87 , 0.90 ) | <0.0001 | 8,176 | 0.88 ( 0.85 , 0.90 ) | <0.0001 | 2,113 | 0.78 ( 0.73 , 0.84 ) | <0.0001 |
|  | 3 | 15,562 | 0.89 ( 0.87 , 0.91 ) | <0.0001 | 8,136 | 0.88 ( 0.85 , 0.91 ) | <0.0001 | 2,097 | 0.78 ( 0.73 , 0.84 ) | <0.0001 |
| IPAQ-PA | 1 | 15,562 | 0.94 ( 0.93 , 0.95 ) | <0.0001 | 8,136 | 0.95 ( 0.93 , 0.97 ) | <0.0001 | 2,097 | 0.95 ( 0.91 , 0.99 ) | 0.0119 |
|  | 2 | 15,562 | 0.98 ( 0.96 , 0.99 ) | 0.00194 | 8,136 | 0.98 ( 0.96 , 1.00 ) | 0.0642 | 2,097 | 0.97 ( 0.93 , 1.01 ) | 0.197 |
|  | 3 | 15,562 | 0.98 ( 0.97 , 1.00 ) | 0.0263 | 8,136 | 0.99 ( 0.97 , 1.01 ) | 0.252 | 2,097 | 0.99 ( 0.94 , 1.03 ) | 0.519 |
| CRF | 1 | 1,393 | 0.67 ( 0.63 , 0.71 ) | <0.0001 | 720 | 0.69 ( 0.64 , 0.75 ) | <0.0001 | 168 | 0.68 ( 0.56 , 0.81 ) | <0.0001 |
|  | 2 | 1,393 | 0.70 ( 0.65 , 0.76 ) | <0.0001 | 720 | 0.76 ( 0.68 , 0.84 ) | <0.0001 | 168 | 0.68 ( 0.54 , 0.85 ) | 0.000614 |
|  | 3 | 1,389 | 0.71 ( 0.65 , 0.76 ) | <0.0001 | 717 | 0.76 ( 0.68 , 0.85 ) | <0.0001 | 167 | 0.70 ( 0.56 , 0.87 ) | 0.00171 |
|  |  | Hemorrhagic stroke |  |  | Heart failure |  |  | Atrial fibrillation |  |  |
|  | Model | N events | HR (95 % CI) | P-value | N events | HR (95 % CI) | P-value | N events | HR (95 % CI) | P-value |
| Grip strength | 1 | 1,012 | 0.91 ( 0.85 , 0.99 ) | 0.0203 | 928 | 0.48 ( 0.45 , 0.52 ) | <0.0001 | 4,449 | 0.80 ( 0.78 , 0.83 ) | <0.0001 |
|  | 2 | 1,012 | 0.88 ( 0.80 , 0.96 ) | 0.00377 | 928 | 0.69 ( 0.63 , 0.75 ) | <0.0001 | 4,449 | 0.96 ( 0.92 , 1.00 ) | 0.0807 |
|  | 3 | 1,009 | 0.89 ( 0.81 , 0.97 ) | 0.00828 | 923 | 0.71 ( 0.64 , 0.78 ) | <0.0001 | 4,423 | 0.96 ( 0.92 , 1.01 ) | 0.085 |
| IPAQ-PA | 1 | 1,009 | 0.93 ( 0.88 , 1.00 ) | 0.0351 | 923 | 0.71 ( 0.66 , 0.76 ) | <0.0001 | 4,423 | 0.97 ( 0.94 , 0.99 ) | 0.0192 |
|  | 2 | 1,009 | 0.93 ( 0.87 , 0.99 ) | 0.0207 | 923 | 0.81 ( 0.76 , 0.86 ) | <0.0001 | 4,423 | 1.02 ( 0.99 , 1.05 ) | 0.237 |
|  | 3 | 1,009 | 0.99 ( 0.94 , 1.03 ) | 0.519 | 923 | 0.83 ( 0.78 , 0.89 ) | <0.0001 | 4,423 | 1.02 ( 0.99 , 1.05 ) | 0.189 |
| CRF | 1 | 81 | 0.98 ( 0.74 , 1.30 ) | 0.881 | 68 | 0.39 ( 0.30 , 0.52 ) | <0.0001 | 428 | 0.64 ( 0.57 , 0.72 ) | <0.0001 |
|  | 2 | 81 | 0.93 ( 0.65 , 1.35 ) | 0.72 | 68 | 0.42 ( 0.31 , 0.58 ) | <0.0001 | 428 | 0.63 ( 0.55 , 0.73 ) | <0.0001 |
|  | 3 | 81 | 0.95 ( 0.66 , 1.38 ) | 0.793 | 68 | 0.44 ( 0.32 , 0.60 ) | <0.0001 | 428 | 0.63 ( 0.55 , 0.72 ) | <0.0001 |
| The effects are in SD-units in fitness and physical activity traits. Model adjustments: 1. Age, gender and region. 2. Age, gender, region, diabetes, smoking, systolic blood pressure, body mass index, lipid medication, height, ethnicity and Townsend index. 3. All in 2) plus IPAQ-PA (grip strength analyses) or grip strength (IPAQ-PA analyses). Analyses of CRF were adjusted for both IPAQ-PA and grip strength.<br>Abbreviations: CVD, cardiovascular disease; CHD, coronary heart disease; IPAQ-PA, physical activity assessed by international physical activity questionnaire; CRF, cardiorespiratory fitness; HR, hazard ratio; CI, confidence interval. |  |  |  |  |  |  |  |  |  |  |

Higher levels of IPAQ-PA were associated with lower risk of CVD events and all-cause death, but the effects were more modest than for grip strength **(Table 2, Table S1).** Associations with the composite CVD outcome, heart failure, and all-cause death remained significant after adjusting for confounding factors and grip strength. Physical activity assessed with a wrist-worn accelerometer had strongest inverse association with all-cause death, when compared to all other measures (HR=0.52, 95% CI 0.46-0.58, **Table S1).**

In a subgroup analysis including 66,652 individuals that underwent a submaximal fitness test, CRF was strongly inversely associated with all CVD events, except hemorrhagic stroke (no. of events=81). When compared with the other two measures of physical activity and fitness, we observed a more pronounced inverse association of CRF with AF (HR, 0.64, 95% CI, 0.57-0.72), which was even stronger when adjusted for other factors, including grip strength and IPAQ-PA.

There were some evidence of nonlinear effects of fitness and physical activity on CVD events **(Figure S2-S4)** and all-cause death **(Figure S5).** In particular, the effect of IPAQ-PA was U-shaped for CVD, CHD, and all-cause death (P_noniinearity_ < 0.0001). However, the association between the objective measurement of physical activity and all-cause death did not support an U-shaped association **(Figure S5).**

### Interactions between fitness, physical activity, and genetic determinants of CHD

Overall, individuals in the highest tertiles of the CHD- and AF-GRSs showed increased risk for incident CHD and AF when compared to those in the lowest tertile (HR, 1.91, 95% CI 1.73-2.10 and HR, 2.08, 95% CI 1.81-2.39, respectively; P for trend < 0.0001). Further adjustment for traditional CVD risk factors (BMI, smoking, lipid medication, systolic blood pressure, and diabetes), grip strength and physical activity did not change the results notably (HR=1.88, 95% CI 1.70-2.07 for CHD and HR=2.08, 95% CI 1.81-2.39 for AF).

Grip strength demonstrated an inverse association with incident CHD and AF in each GRS group, although the effect was strongest in the lowest GRS group (P_interaction_ = 0.006 for CHD, P_interaction_ = 0.03 for AF, Figure 1). For was similar for CHD, but not for AF (P_interaction_ = 0.02 and P_interaction_ = 0.35, respectively, **Figure S6).** For IPAQ-PA, the associations between CHD were modest and no significant differences by GRS groups were observed (P_interaction_ = 0.08 for CHD, P_interaction_ = 0.50 for AF, **Figure S7).**

**Figure 1.**
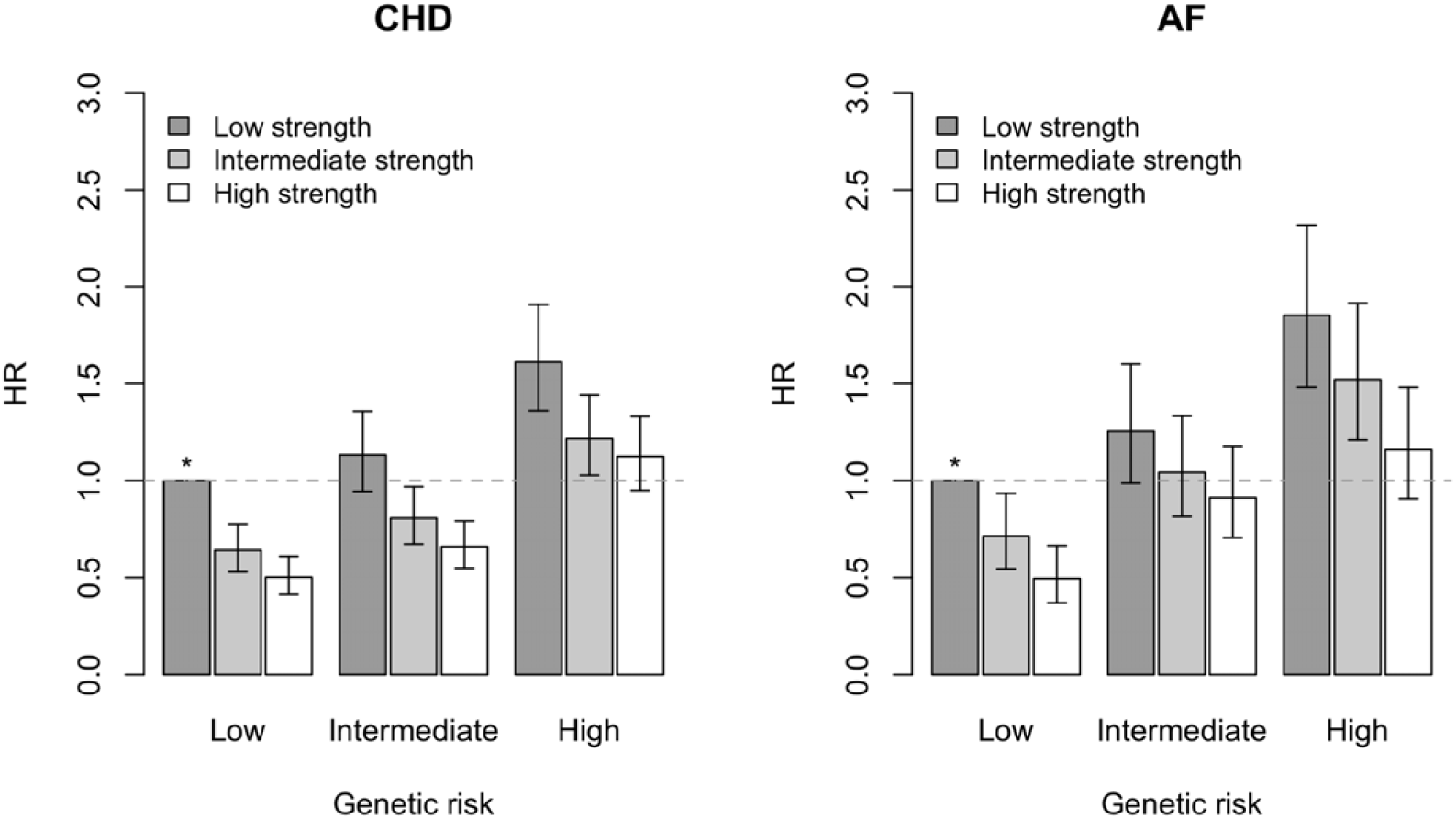
Hazard ratios with 95% confidence intervals for coronary heart disease (CHD) and atrial fibrillation (AF) according to tertiles of genetic risk and grip strength. * Reference group.

### Genome-wide association studies of measures of fitness and physical activity

In our discovery analysis, we found 14 genome-wide significant independent loci associated with grip strength, of which six was formally replicated (alpha: 0.05/14, Table 3, Figure S8). The strongest signal was in chromosome 16 in the *FTO* locus (rs28429148, P=7.4×10^-21^). Our lead SNP is in high LD (R^2^>0.8) with the other *FTO* variants previously associated with cardio-metabolic traits including obesity^21,22^ and lipids^23^. The other replicated SNPs were located in or near *TFAP2B* (P=1.1×10^-11^), *TMEM18/FAM150B* (two independent loci, P=9.8×10^-11^ and P=7.4×10^-10^), *RAPSN* (P=2.2×10^-9^), and *ADCY3* (P=4.2×10^-8^). Variants in these regions have been previously associated with several traits including height and obesity **(Figures S9-S13).** None of the loci reached genome-wide significance for IPAQ-PA.

**Table 3.** Genetic loci associated with grip strength in discovery and replication datasets.

|  |  |  |  |  |  |  |  |  |  |  |  |  |
| --- | --- | --- | --- | --- | --- | --- | --- | --- | --- | --- | --- | --- |
|  |  |  |  |  |  |  | *Discovery* |  |  | *Replication* |  |  |
| SNP | CHR | Position | Gene | EA | OA | EAF | Beta | SE | P-value | Beta | SE | P-value |
| rs28429148 | 16 | 53798319 | *FTO* | G | A | 0.5667 | 0.04 | 0.004 | $7.4 \times 10^{-21}$ | 0.023 | 0.006 | $1.6 \times 10^{-4}$ |
| rs72892910 | 6 | 50816887 | *TFAP2B* | G | T | 0.8285 | 0.038 | 0.005 | $1.1 \times 10^{-11}$ | 0.024 | 0.008 | 0.002 |
| rs12623218 | 2 | 632146 | *TMEM18* | T | A | 0.1724 | 0.036 | 0.005 | $9.8 \times 10^{-11}$ | 0.028 | 0.008 | $3.5 \times 10^{-4}$ |
| rs62107261 | 2 | 422144 | *FAM150B* | T | C | 0.9517 | -0.06 | 0.009 | $7.4 \times 10^{-10}$ | -0.057 | 0.014 | $4.1 \times 10^{-5}$ |
| rs12361415 | 11 | 47474146 | *RAPSN* | T | G | 0.7131 | -0.027 | 0.005 | $2.2 \times 10^{-9}$ | -0.022 | 0.006 | $6.0 \times 10^{-4}$ |
| rs556981345 | 2 | 25115122 | *ADCY3* | C | CT | 0.5626 | 0.024 | 0.004 | $4.2 \times 10^{-8}$ | 0.027 | 0.006 | $8.0 \times 10^{-6}$ |
| Effects are in SD-units of grip strength. |  |  |  |  |  |  |  |  |  |  |  |  |
| Abbreviations: SNP, single-nucleotide polymorphism; CHR, chromosome; EA, effect allele, OA, other allele; EAF, effect allele frequency; SE, standard error. |  |  |  |  |  |  |  |  |  |  |  |  |

In our pooled analysis of all samples **(Table S2),** 27 loci reached genome-wide significance for grip strength, and the strongest association was observed for another SNP in *FTO* (rs56094641, P=3.8×10^-24^). Further adjustment for height did not change the effect estimates of the 27 grip strength loci (Pearson correlation=1.0 between beta coefficients). For IPAQ-PA, the pooled analysis yielded two loci (rs9316077 in *SMIM2,* P=1.4×10^-8^ and rs146370962 in *PSAT1,* P=3.2×10^-8^). We observed inflation in test statistics (lambda=1.16 for grip strength and 1.06 for IPAQ-PA), which is expected under polygenic inheritance in large samples^24^.

For DEPICT analyses, we included 200 grip strength and 51 IPAQ-PA variants with at a lower level of evidence (P<10^-5^) from our pooled GWAS (see list of SNPs in **Table S3)** to increase power. DEPICT analyses identified cardiovascular and musculoskeletal tissue as the most significant physiological system for grip strength. However, none of the DEPICT analyses were formally significant (FDR<0.05, **Figures S14-S15).** GTEx analysis identified several genes significantly regulated by the grip strength SNPs **(Table S4).** For example, one of the replicated lead variants, rs72892910 near *TFAP2B,* was a significant eQTL for *TFAP2B* in lung. Another of the replicated lead variants, rs12361415 near *RAPSN,* had evidence of several significant eQTLs (*C1QTNF4, MADD, RAPSN).* This is a particularly interesting locus, as the mutations in *RAPSN* are known to cause congenital myasthenic syndrome, characterized by muscle weakness^25,26^.

The allelic sum of grip strength loci was associated with CRF, IPAQ-PA and accelerometry, whereas IPAQ-PA loci showed association with physical activity measured with accelerometer **(Table S5).**

### Mendelian randomization

From our 27 grip strength loci, 20 lead variants and two proxies were found in the CARDIoGRAMplusC4D summary data. The F-statistics for GRSs using 22 grip strength and two IPAQ-PA variants were 688 and 40, respectively. At an alpha level 0.05, the statistical power to detect causal estimates for grip strength was 95.6% with the observed effect size of 0.79 (per SD-increase of grip strength, see Table 2). For IPAQ-PA, the power was 3.9% (effect size 0.95, Table 2). Consequently, we decided to proceed with MR analyses for grip strength, but not with IPAQ-PA.

The results from the two-sample MR indicated a causal effect of grip strength on CHD (IVW: OR = 0.56, 95% CI 0.44-0.70, P=1.4×10^-7^; Weighted median: OR = 0.69, 95% CI 0.51-0.95, P=0.01, **Table S6).** The MR Egger results for were non-significant, but the effect estimate was consistent with the other methods (OR = 0.51, 95% CI 0.19-1.36, P=0.17). Our analyses indicated no significant heterogeneity or directional horizontal pleiotropy (all P>0.10; and no evidence upon visual inspection, Figure 2). Hierarchical clustering using Euclidean distance and Ward method^27-28^ separated SNPs into three clusters; BMI-associated SNPS (group 1), others (group 2), and FTO (group 3) **(Figure S16).** For eight variants, the association with BMI was stronger than for grip strength. None of these variants were included in the group 2. The causal effects were similar when performing MR for SNPs in group 1 (IVW: OR = 0.59,95% CI 0.43-0.81, P=0.0005) and 2 (IVW: OR = 0.56, 95% CI 0.37-0.84, P=0.003) separately, suggesting that the effect grip strength on CHD is mediated through obesity, but also through some other unknown pathway.

**Figure 2.**
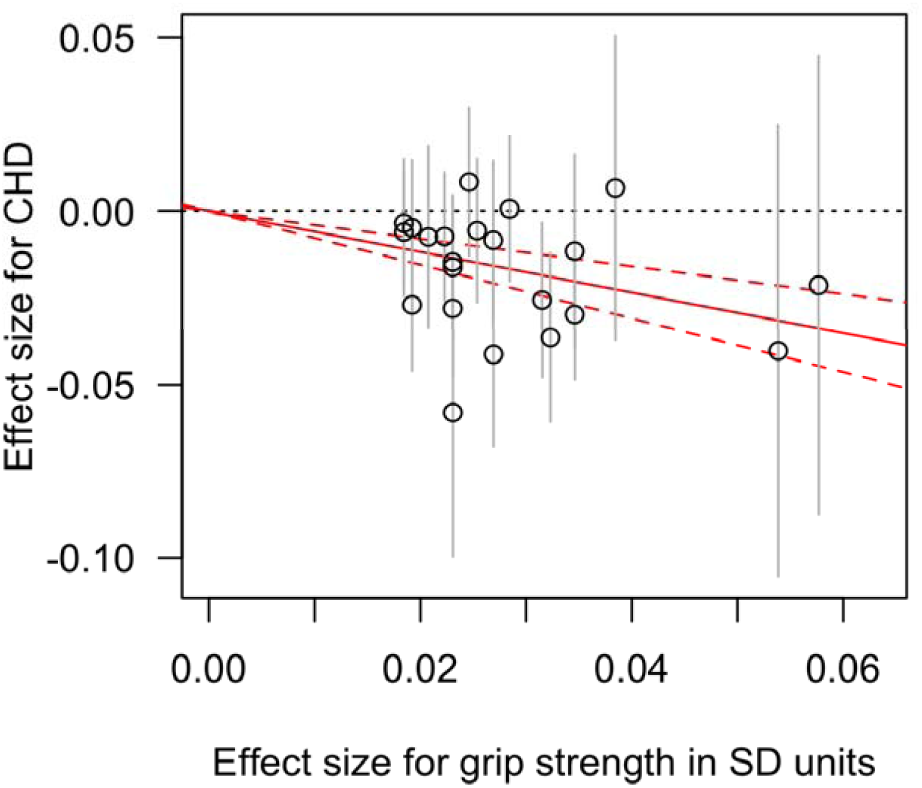
Effects of genetic variants on grip strength and risk for coronary heart disease including 22 variants with genome-wide significant association with grip strength in the pooled analysis of discovery and replication samples.

## Discussion

### Principal Findings

In this study of up to 502,635 individuals from the general population, we report the associations of objective and subjective measures of fitness and physical activity with six cardiovascular outcomes and total mortality; explore the role of gene-environment interactions in CHD development; establish genetic determinants of fitness and physical activity; and address the causal role of grip strength in CHD development. Our main findings are several-fold. First, in an observational study of unprecedented size, we establish that grip strength, physical activity and CRF have strong inverse associations with different types of incident CVD events and all-cause death, and that accelerometry-based physical activity is associated with all-cause death. Second, we show significant interaction effects between fitness and genetic risk, with stronger associations of grip strength with CHD and AF in individuals with lowest genetic risk. Third, we report 27 loci (6 with formal replication), associated with grip strength and 2 loci with IPAQ-PA providing new leads regarding biological processes involved in individual variation in and response to fitness and physical activity. Fourth, our analyses indicate that grip strength is causally related to CHD, which highlights the importance of strength in maintaining good cardiovascular health.

### Comparison with Prior Literature

Studies using objective measurements of fitness and physical activity (such as VO_2_) involve intensive and time-consuming exercise tests, and are rarely implemented in large samples. Two notable exceptions utilized a large database of exercise testing in Swedish young men performed at military service conscription^2,29^. As in our study, these studies reported inverse associations of grip strength and CRF with CVD. In contrast to our results, Andersen et al.^2^ reported a positive association between CRF and AF. However, the effect was clearly attenuated when adjusting for available confounders, the availability of which was limited compared to our study. Moreover, the study was conducted in young men, so the results may not be generalizable to general population, while our analyses were performed in middle-aged to elderly men and women. Indeed, it has been suggested that the exercise-associated AF affects mostly male endurance athletes and the etiology might be different from that of general AF^30^, presumably examined in the present study. In the present study, we were able to adjust for a large set of potential confounders, and this strengthened the inverse association of CRF with AF.

Due to the challenges of measuring fitness and physical activity, studies relating these traits with prospective CVD in the general population have previously been limited by small sample size or lack of measurement accuracy. Attempts to combine data in meta-analyses have had to use broad categories of fitness and physical activity to harmonize the data across many small studies^31,32^. Compared to these studies, the UK Biobank has a clear advantage as the traits were measured in the same way in over 500,000 individuals. Nevertheless, our results are consistent with previous meta-analyses reporting weaker associations for questionnaire-based physical activity assessment compared to more objective measures^31^. This finding suggests that associations of physical activity with outcome are most likely underestimated in studies using questionnaire data. Moreover, our results showing an U-shaped association between self-reported, but not objective, physical activity and all-cause death, suggests that some additional factors might explain an increased risk in those individuals reporting very high values of physical activity.

Prior genetic studies of fitness and physical activity have suffered insufficient statistical power leading to contradictory results^33^. To our knowledge, this is the largest study to date to combine large-scale physical activity and fitness data with genetic information. This resulted in the discovery of a much large number of genetic variants associated with grip strength compared to a recent study^34^.

### Biological Insights Gained from Genetic Analyses

We report 29 novel loci associated with fitness or physical activity. We detected several significant eQTLs in tissues relevant for CVD, which could help highlighting causal genes and to understand the biological mechanisms underlying protective effects of fitness in CVD. Our large gene-environment interaction analyses of CHD and AF indicated that maintaining good strength can compensate for genetic risk of these diseases, as the inverse associations of grip strength and CRF with disease was seen in each category of genetic risk. The effect was strongest in the lowest genetic risk group, suggesting that individuals with elevated genetic risk for cardiovascular events need to maintain higher fitness level to gain similar risk reductions than those without genetic burden.

### Clinical Implications

Based on our two-sample MR analysis, grip strength seems causally related to CHD, suggesting that maintaining strength is likely to help preventing future disease events. This is in line with a recent smaller Mendelian randomization study of grip strength^35^ and previous randomized controlled trials showing favorable effects of resistance training on cardiovascular risk factors^36^. Future studies evaluating causality of strength versus aerobic training on subclinical or clinical cardiovascular outcomes could help to tailor exercise programs for individuals with elevated risk for these diseases.

Our results showing the inverse association of grip strength also in genetically predisposed individuals points to potential advantages of genetic risk profiling for better detection of individuals at risk for CVD. Even though more information is needed to evaluate how people understand the genetic risks, the knowledge that lifestyle choices have substantial effects on the disease risks could encourage individuals to initiate a healthier lifestyle to reduce their overall risk.

### Strengths and Limitations

Strengths of this study include that this is the largest study to date to combine objective and subjective measures of fitness and physical activity, genome-wide array data and prospective follow-up of CVD events. Further, we have used state-of-the-art methods including a conservative analytical framework with strict multiple testing correction, replication of genetic findings and robust analysis methods for MR analyses.

Our study also has several limitations. First, although this was the largest GWAS ever performed for grip strength and physical activity, the sample size was not large enough to discover sufficient number of genetic variants for physical activity and CRF to perform adequately powered MR. Second, in our MR analysis, we used multiple SNPs in the IV, which may increase the risk of including pleiotropic effects. Also, given the complex nature of grip strength^37^, we can have included genetic variants that are proxies of some other biological phenomena rather than strength. That said, our comprehensive analytical framework did not indicate any presence of horizontal pleiotropy, which would violate MR assumptions, while there may still be vertical pleiotropy (or mediation) that does not violate the assumptions. Further, grip strength measures mainly upper body strength. However, it is highly correlated with knee extension muscle strength (r= 0.772 to 0.805)^38^, and also, our results show that genetic variants for grip strength are also associated with CRF and physical activity. Thus, grip strength seems to be a good indicator of overall fitness. Finally, although our analyses were conducted in a cohort with different ethnicities, the majority of participants were of European ancestry. Specifically, our GWAS were conducted in European samples only to avoid population stratification. Hence, the generalizability of the results to other ethnicities is not fully understood.

### Conclusions

In conclusion, different measures of fitness and physical activity demonstrated strong inverse associations with future CVD events and all-cause death in a large population-based sample. In particular, grip strength was causally related with CHD, highlighting the importance of strength in prevention of cardiovascular events. Maintaining good strength can also compensate for genetic risk of CHD and AF.

## Acknowledgements

This research has been conducted using the UK Biobank Resource under Application Number 13721. Data on coronary heart disease have been contributed by CARDIoGRAMplusC4D investigators and have been downloaded from <u>www.cardiogramplusc4d.org</u>. The research was performed with support from National Institutes of Health (1R01HL135313-01; 1R01DK106236-01A1) and Knut and Alice Wallenberg Foundation (2013.0126). ET was supported by the Finnish Cultural Foundation.

## Declaration of interests

Erik Ingelsson is a scientific advisor for Precision Wellness, Cellink and Olink Proteomics for work unrelated to the present project.

## Authors' contributions

Design of the study: ET, EI; Analysis and interpretation of data: ET, SG, EI; Drafting the manuscript: ET; Critical revision of the manuscript: all authors; Final approval of the version to be published: all authors.

